# Numb-associated kinases are required for SARS-CoV-2 infection and are cellular targets for therapy

**DOI:** 10.1101/2022.03.18.484178

**Authors:** Marwah Karim, Sirle Saul, Luca Ghita, Malaya Kumar Sahoo, Chengjin Ye, Nishank Bhalla, Jing Jin, Jun-Gyu Park, Belén Martinez-Gualda, Michael Patrick East, Gary L. Johnson, Benjamin A. Pinsky, Luis Martinez-Sobrido, Christopher R. M. Asquith, Aarthi Narayanan, Steven De Jonghe, Shirit Einav

## Abstract

The coronavirus disease 2019 (COVID-19) pandemic caused by the severe acute respiratory syndrome coronavirus 2 (SARS-CoV-2) continues to pose serious threats to global health. We previously reported that AAK1, BIKE and GAK, members of the Numb-associated kinase family, control intracellular trafficking of multiple RNA viruses during viral entry and assembly/egress. Here, using both genetic and pharmacological approaches, we probe the functional relevance of NAKs for SARS-CoV-2 infection. siRNA-mediated depletion of AAK1, BIKE, GAK, and STK16, the fourth member of the NAK family, suppressed SARS-CoV-2 infection in human lung epithelial cells. Both known and novel small molecules with potent AAK1/BIKE, GAK or STK16 activity suppressed SARS-CoV-2 infection. Moreover, combination treatment with the approved anti-cancer drugs, sunitinib and erlotinib, with potent anti-AAK1/BIKE and GAK activity, respectively, demonstrated synergistic effect against SARS-CoV-2 infection *in vitro*. Time-of-addition experiments revealed that pharmacological inhibition of AAK1 and BIKE suppressed viral entry as well as late stages of the SARS-CoV-2 life cycle. Lastly, suppression of NAKs expression by siRNAs inhibited entry of both wild type and SARS-CoV-2 pseudovirus. These findings provide insight into the roles of NAKs in SARS-CoV-2 infection and establish a proof-of-principle that pharmacological inhibition of NAKs can be potentially used as a host-targeted approach to treat SARS-CoV-2 with potential implications to other coronaviruses.

## 1. Introduction

SARS-CoV-2 responsible for the COVID-19 pandemic continues to pose a major global health challenge (Zhou et al., 2020; Mallah et al., 2021). While effective vaccines are available to reduce COVID-19 severity, limited access to vaccines in various parts of the world, vaccine hesitancy, and continuous emergence of new viral variants are among the ongoing challenges. A few direct-acting antivirals (DAAs), including the protease inhibitor nirmatrelvir (Owen et al., 2021) and the polymerase inhibitor molnupiravir (Jayk Bernal et al., 2022), have demonstrated promising reduction of hospitalization and death rates. Nevertheless, widespread use of these DAAs may lead to the emergence of resistant viral variants. Indeed, viral escape mutants were identified in cultured cells infected with various coronaviruses under treatment with the polymerase inhibitors molnupiravir (Agostini et al., 2019) and remdesivir (Stevens et al., 2022). There is thus a need for host-targeted antiviral drugs that will effectively suppress viral replication with a high barrier to resistance.

SARS-CoV-2 entry into target cells is mediated by the angiotensin-converting enzyme 2 (ACE2) receptor and is enhanced by neuropilin-1 (NRP-1) (Cantuti-Castelvetri et al., 2020). Fusion of the SARS-CoV-2 envelope is thought to occur primarily at the plasma membrane where the cellular transmembrane protease serine 2 (TMPRSS2) cleaves the viral spike (S) protein at the S2’ site following its pre-cleavage at the S1/S2 site (Hoffmann et al., 2020). Nevertheless, fusion with endosomal membranes following clathrin-mediated endocytosis (CME) and cleavage of the S protein by cathepsin L has also been reported, particularly in cells expressing low level of TMPRSS2 (Bayati et al., 2021). Regardless of the entry mechanism, the replicase complex is then translated from the RNA genome followed by viral RNA replication. Virions are thought to assemble in the endoplasmic reticulum-golgi intermediate compartment (ERGIC) and egress via the secretory pathway (Klein et al., 2020).

The Numb-associated kinases family of Ser/Thr kinases is composed of adaptor-associated kinase 1 (AAK1), BMP-2 inducible kinase (BIKE/BMP2K), cyclin G-associated kinase (GAK), and serine/threonine kinase 16 (STK16). These kinases share only limited homology in their kinase domain and low homology in other protein regions (Sorrell et al., 2016). NAKs have been shown to regulate intracellular membrane trafficking (Sorensen and Conner, 2008; Sato et al., 2009). AAK1 and GAK phosphorylate the endocytic adaptor protein complex 2 (AP2M1) and the secretory AP complex 1 (AP1M1), thereby stimulating their binding to cellular cargo (Conner and Schmid, 2002). BIKE was identified as an accessory protein on a subset of clathrin-coated vesicles that binds the endocytic adaptor NUMB and phosphorylates AP2M1 (Kearns et al., 2001; Krieger et al., 2013; Sorrell et al., 2016; Pu et al., 2020). STK16, the most distantly related member of the NAK family, regulates various physiological activities including Golgi assembly, trans-Golgi network (TGN) protein secretion and sorting as well as cell cycle and TGF-beta signaling (In et al., 2014; López-Coral et al., 2018).

We have previously discovered that AAK1 and GAK mediate intracellular trafficking of hepatitis C virus (HCV), dengue virus (DENV) and Ebola virus (EBOV) during viral entry and assembly/egress (Neveu et al., 2012; Neveu et al., 2015; Bekerman et al., 2017; Xiao et al., 2018). We have recently shown a requirement for BIKE in DENV infection (Pu et al., 2020), beyond its reported role in human immunodeficiency virus-1 (HIV-1) infection (Zhou et al., 2008). Additionally, we have demonstrated that pharmacological inhibition of AAK1/BIKE and/or GAK activity by known or novel small molecules has a broad-spectrum antiviral potential both *in vitro* and *in vivo* (Bekerman et al., 2017; Pu et al., 2020).

While AAK1 was proposed as a cellular target of SARS-CoV-2 based on *in silico* analysis (Richardson et al., 2020; Stebbing et al., 2020), the roles of NAKs in SARS-CoV-2 infection and as candidate targets for anti-SARS-CoV-2 therapy remain largely uncharacterized. Here, we sought to probe the functional relevance of the four members of the NAK family in SARS-CoV-2 infection and determine their role as candidate anti-SARS-CoV-2 targets.

## 2. Materials and Methods

### 2.1. Cells

Calu-3 cells (ATCC) were grown in medium (DMEM; Gibco) supplemented with 10% fetal calf serum (FCS; Omega Scientific Inc.), 1% Pen-strep (Gibco) and 1% nonessential amino acids (NEAA, Gibco). The African green monkey kidney cell line (Vero E6, ATCC) was maintained in DMEM supplemented with 10% FCS, 1%L-glutamine 200mM, 1% Pen-strep, 1% NEAA, 1% HEPES (Gibco), 1% Sodium pyruvate (Thermofisher scientific). Vero E6-TMPRSS2 (JCRB cell bank, #cat JCRB1819) were maintained in DMEM supplemented with 10% FCS, 1% Pen-strep and 1mg/ml G418 (Gibco, cat. #10131035). HEK-293T (ATCC) cells were grown in DMEM supplemented with 10% FCS, 1%L-glutamine, and 1% Pen-strep. Cells were maintained in a humidified incubator with 5% CO_2_ at 37°C and were tested negative for mycoplasma.

### 2.2. Compounds

The following reagents were used: (5Z)-7-oxozeaenol (Cayman Chemical), sunitinib malate, (Selleckchem), erlotinib (LC Laboratories), gefitinib (Selleckchem), baricitinib (a gift from Dr. Chris Liang), SGC-GAK-1 and STK16-IN-1 (MedChemExpress). RMC-76 (compound 21b) and RMC-242 (compound 7b) were prepared as previously described (Verdonck et al., 2019; Martinez-Gualda et al., 2021).

### 2.3. Viral stocks preparation and sequencing

The 2019-nCoV/USA-WA1/2020 SARS-CoV-2 isolate (NR-52281) (BEI Resources) was passaged 3 times in Vero E6-TMPRSS2 cells. The rSARS-CoV-2/WT and rSARS-CoV-2 expressing Nluc-reporter genes (rSARS-CoV-2/Nluc) were generated as previously described (Chiem et al., 2020; Ye et al., 2020). Briefly, Vero E6 cells were transfected using Lipofectamine 2000 (Thermo Fisher) with 4 mg/well of pBeloBAC11-SARS-CoV-2/WT or – Nluc-2A. At 12 hours post-transfection, medium was replaced by DMEM with 2% FBS. At 72 hours, P0 virus-containing tissue culture supernatants were collected and stored at −80°C. Following titration, P0 virus stock was used to generate a P1 stock by infecting Vero E6 monolayers with multiplicity of infection (MOI) of 0.0001 for 72 hours. P1 virus was passaged twice in Vero E6-TMPRSS2 cells. Viral titers were determined by standard plaque assay on Vero E6 cells.

Work involving WT SARS-CoV-2 was conducted at the BSL3 facilities of Stanford University, George Mason University, and Texas Biomedical Research Institute according to CDC and institutional guidelines. SARS-CoV-2 stocks were deep sequenced on a MiSeq platform (Illumina). SARS-CoV-2 whole-genome amplicon-based sequencing was conducted by adapting an existing whole genome sequencing pipeline for poliovirus genotyping as described (Wang et al., 2021). All the experiments were performed using a P3 SARS-CoV-2 USA-WA1/2020, rSARS-CoV-2/WT or rSARS-CoV-2/Nluc containing 100% WT population with no deletion in the Spike multi-basic cleavage site.

### 2.4. rVSV-SARS-CoV-2-S pseudo-virus production

As described in (Saul et al., 2021), HEK-293T cells were transfected with 30 μg of a Spike expression plasmid. After 24 hours, the medium was replaced and cells were treated with valproic acid (VPA). Following a 4 hour incubation, the cells were infected with VSV-G pseudo typed ΔG-luciferase VSV virus (MOI=3). At 6 hours post-infection (hpi), cells were washed with PBS, and fresh medium containing anti-VSV-G hybridoma was added to neutralize the residual VSV-G pseudo-virus. Twenty-four hours later, culture supernatant was harvested, clarified by centrifugation, filtered (0.22 μm) and stored at −80°C. rVSV-SARS-CoV-2-S was titrated on Vero cells via luciferase assay, and TCID_50_ was determined. Positive wells were defined as having luminescence values at least 10-fold higher than the cell background.

### 2.5. Plaque assay and nano-luciferase (Nluc) assay

For plaque assays, monolayers of Vero E6 cells were infected with WT SARS-CoV-2 or rSARS-CoV-2/Nluc for 1 hour at 37 °C. Cells were then overlaid with MEM and carboxymethyl cellulose and incubated at 37 °C. At 72 hpi, the overlay was removed, and the cells were submerged in 70% ethanol for fixation and viral inactivation followed by crystal violet staining. Nluc expression was determined on a plate reader GloMax Discover Microplate Reader (Promega) at 24 hpi in the infected cell supernatants using a luciferin solution obtained from the hydrolysis of its O-acetylated precursor, hikarazine-103 (prepared by Dr. Yves Janin, Pasteur Institute, France) as a substrate (Coutant et al., 2019; Coutant et al., 2020).

### 2.6. Time-of-addition experiment

Calu-3 cells were infected with rSARS-CoV-2/WT (MOI=1). Two hpi, the virus was removed and cells were washed twice with PBS. At 2, 5 and 8 hpi, 5 μM of RMC-76 or 0.1% DMSO were added. Cell culture supernatants were collected at 10 hpi, and infectious viral titers were measured by plaque assay.

### 2.7. RNA interference

The following siRNAs (10 pmole) were transfected into Calu-3 cells using Dharmafect-4 transfection reagent (#cat: T-2004-02, Dharmacon) 48 hours prior to infection with SARS-CoV-2: siAAK1 (ID: s22494, Ambion), siBIKE (M-005071-01, Dharmacon), siGAK (ID: s5529, Ambion), siSTK16 (AM51331, Life Technology) and siNT (D-001206-13-05, Dharmacon).

### 2.8. Infection assays

Calu-3 and Vero E6 cells were infected with SARS-CoV-2 (USA-WA1/2020) or rSARS-CoV-2/Nluc in triplicates (MOI=0.05) in DMEM containing 2% FCS. After 1-hour incubation at 37 °C, the viral inoculum was removed, cells were washed and new medium was added. Culture supernatants were harvested at 24 hpi and the viral titer was measured via plaque or Nluc assays.

### 2.9. Entry assays

Calu-3 cells were infected with WT rSARS-CoV-2/WT (MOI=1) or a high inoculum of rVSV-GP SARS-CoV-2. Following 2-hour incubation, the viral inoculum was removed, cells were washed with PBS and lysed in TRIzol LS, and viral RNA level measured by RT-qPCR.

### 2.10. RT-qPCR

RNA was extracted from Calu-3 lysates using Direct-zol RNA Miniprep Plus Kit (Zymo Research) and reverse transcribed using High-Capacity cDNA RT kit (Applied Biosystems) according to the manufacturer’s instructions. Primers and PowerUp SYBR Green Master Mix (Applied Biosystems) were added to the samples, and PCR reactions were performed with QuantStudio3 (Applied Biosystems) in triplicates. Target genes were normalized to the housekeeping gene (GAPDH). Sequences of primers used for RT-qPCR are available upon request.

### 2.11. Pharmacological inhibition

The inhibitors or DMSO were administered to cells 1 hour prior to viral inoculation and maintained for the duration of the experiment. Viral infection was measured via Nluc (rSARS-CoV-2/Nluc) or plaque (SARS-CoV-2) assays.

### 2.12. Viability assays

Cell viability was assessed using alamarBlue reagent (Invitrogen) according to the manufacturer’s protocol. Fluorescence was detected at 560 nm on an InfiniteM1000 plate reader (Tecan) and GloMax Discover Microplate Reader (Promega).

### 2.13. NanoBRET assay

NanoBRET for RMC-242 was performed at Carna Biosciences. Briefly, HEK293 cells were transiently transfected with the NanoLuc^®^ Fusion DNA and incubated at 37°C. 20 hours post-transfection, NanoBRET^™^ tracer reagent and RMC-242 were added to the cells and incubated at 37°C for 2 hours. Nanoluciferase-based bioluminescence resonance energy transfer (BRET) was measured using NanoBRET^™^ Nano-Glo^®^ Substrate on a GloMax^®^ Discover Multimode Microplate Reader (Promega).

### 2.14. *In vitro* kinase assays

*In vitro* kinase assays to determine IC_50_ of gefitinib for AAK1 and BIKE were performed by the LabChip platform (Nanosyn).

### 2.15. Multiplexed Inhibitor Bead (MIB) affinity chromatography/MS analysis

Kinome profiling was performed as previously described (Duncan et al., 2012; Asquith et al., 2018). Briefly, SUM159 cell lysates were incubated with DMSO or the indicated concentration of RMC-76 or DMSO for 30 min on ice. Kinases fragments were then detected and analyzed by mass spectrometry. Kinase abundance was quantified label-free using MaxQuant software.

### 2.16. Data analysis of combination drug treatment

The MacSynergy II program was used to perform synergy/antagonism analysis, as previously described (Prichard and Shipman, 1996; Neveu et al., 2015; Bekerman et al., 2017). The combination’s effect is determined by subtracting the theoretical additive values from the experimental values. A synergy peak is depicted by the three-dimensional differential surface plot above a theoretical additive plane and antagonism as depressions below it (Prichard and Shipman, 1990). Matrix data sets in 3 replicates were assessed at the 95% confidence level for each experiment. Synergy and log volume were calculated.

### 2.17. Statistical Analysis

All data were analyzed with GraphPad Prism software. Half-maximal effective concentrations (EC50) and half-maximal cytotoxic concentrations (CC50) were measured by fitting of data to a 3-parameter logistic curve. *P* values were calculated by ordinary 1-way ANOVA with Dunnett’s multiple comparisons tests as specified in each figure legend.

## 3. Results

### 3.1. NAKs are required for wild type (WT) SARS-CoV-2 infection

To probe the functional relevance of NAKs to WT SARS-CoV-2 infection, we first used an RNAi approach. Calu-3 (human lung epithelial) cells were depleted of AAK1, BIKE, GAK and STK16 individually via siRNAs (**Figure 1A**), and knock-down efficiency was confirmed via RT-qPCR (**Figure 1B)**. NAKs depletion had no apparent effect on Calu-3 cell viability, as measured by an alamarBlue assay (**Figure 1B**). Nevertheless, it reduced SARS-CoV-2 infection by 89%±1.1% (siAAK1), 91%±5.7% (siBIKE), 91%±5.0% (siGAK), and 96%±4.3% (siSTK16) relative to the non-targeting (siNT) control, as measured via Nluc assay at 24 hours post-infection (hpi) with rSARS-CoV-2/Nluc virus (SARS-CoV-2 USA-WA1/2020 isolate (NR-52281) expressing a nanoluciferase reporter gene)) (Chiem et al., 2020; Ye et al., 2020) (**Figure 1C**). Moreover, a 2-3 log reduction in SARS-CoV-2 titer was measured in the infected cells relative to the siNT control via plaque assay at 24 hpi (**Figure 1D**). These loss-of-function experiments provide genetic evidence that the four NAKs are required for effective SARS-CoV-2 infection.

**Figure 1.**
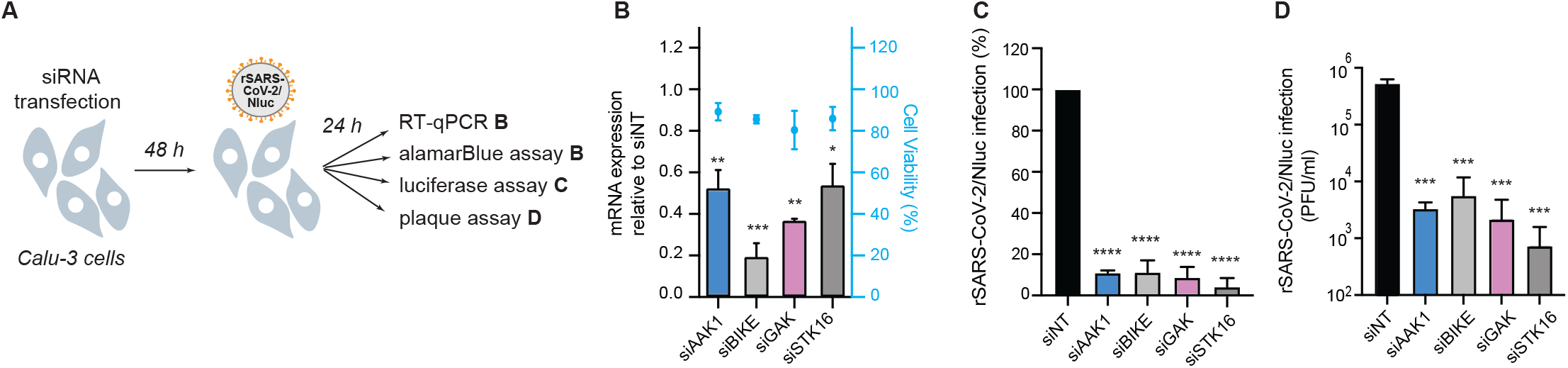
NAKs are required for SARS-CoV-2 infection. (**A**) Schematic of the experiments shown in panels **B-D**. (**B**) Confirmation of siRNA-mediated gene expression knockdown and cell viability in Calu-3 cells transfected with the indicated siRNAs. Shown is gene expression normalized to GAPDH and expressed relative to the respective gene level in the non-target (siNT) control at 48 hours post-transfection measured by RT-qPCR and cell viability measured by alamarBlue assay. (**C**) WT SARS-CoV-2 infection measured at 24 hours post-infection (hpi) of the indicated NAK-depleted Calu-3 cells with rSARS-CoV-2/Nluc (USA-WA1/2020 strain; MOI=0.05) by Nluc assays. (**D**) Viral titers measured at 24 hpi of the indicated NAK-depleted Calu-3 cells with rSARS-CoV-2/Nluc (USA-WA1/2020 strain; MOI=0.05) by plaque assays. Data in all panels are representative of 2 or more independent experiments. Individual experiments had 3 biological replicates, means ± standard deviation (SD) are shown. Data are relative to siNT (**C, D**). *P < 0.05; **P < 0.01; ***P < 0.001; ****P < 0.0001 by 1-way ANOVA followed by Dunnett’s multiple comparisons test. PFU, plaque-forming units.

### 3.2. Known pharmacological inhibitors with anti-NAK activity suppress SARS-CoV-2 infection

To determine whether a similar effect on SARS-CoV-2 infection to that achieved genetically can be achieved pharmacologically, we determined the anti-SARS-CoV-2 activity of a set of 5 compounds: (5Z)-7-oxozeanol, sunitinib, erlotinib, gefitinib and baricitinib (**Table 1 and Figure 2A**), via two orthogonal assays in two distinct cell lines. Vero E6 and Calu-3 cells were pretreated for 1 hour with the compounds followed by infection with WT SARS-CoV-2 (2019-nCoV/USA-WA1/2020) or rSARS-CoV-2/Nluc (Chiem et al., 2021), respectively, both at an MOI of 0.05. The antiviral effect of the compounds was measured at 24 hpi via plaque (Vero E6 cells) or Nluc (Calu-3 cells) assays. Their effect on cellular viability was measured in the infected cells via alamarBlue assays.

**Figure 2.**
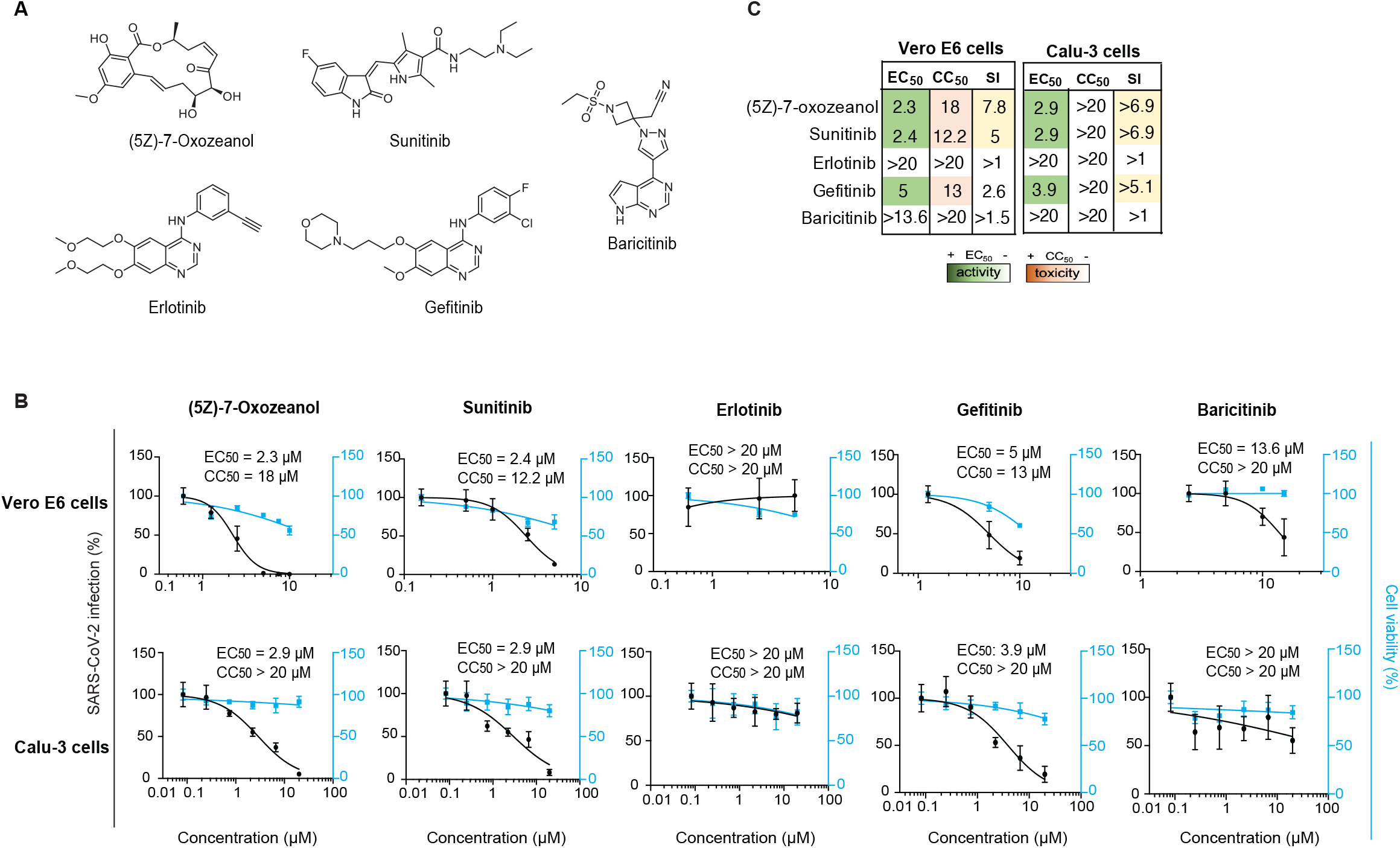
Known compounds with anti-NAK activity suppress SARS-CoV-2 infection. (**A**) Chemical structures of known compounds with anti-NAK activity. (**B**) Dose response of WT SARS-CoV-2 infection to the indicated compounds in Vero E6 cells infected with rSARS-CoV-2/Nluc and in Calu-3 cells infected with USA-WA1/2020 strain (MOIs=0.05) (black) measured via plaque and Nluc assays at 24 hpi, respectively. Dose response of cellular viability to the indicated compounds measured via alamarBlue assays are shown in blue. (**C**) Heat maps of the EC_50_ and CC_50_ values of the inhibitors color-coded based on the antiviral activity and cell viability measured in the indicated cell lines. Selectivity indices (SI, CC_50_ to EC_50_ ratios) greater than 5 are depicted in yellow. Data in all panels are representative of 2 or more independent experiments. Individual experiments had 3 or greater biological replicates. Shown are means ± SD.

**Table 1.** Kinase inhibitors with anti-NAK activity used in this study. Dissociation constant (K_d_), enzymatic activity (IC_50_), percent binding of control (% of control) values measured *in vitro* on the four NAKs and NanoBRET GAK IC_50_ values measured in cells of the indicated kinase inhibitors. The source of kinomeScan data (if available) and other targets of these compounds are also shown. K_d_, IC_50_, and % of control are shown as *, ** and *** respectively. ^ represents data obtained in this study.

| Compound | Status | K<sub>d</sub><sup>*</sup>, IC<sub>50</sub> (nM)<sup>**</sup> or % of control<sup>***</sup> |  |  |  | NanoBRET<br>GAK IC<sub>50</sub> (nM) | Reference | Kinome | Other targets |
| --- | --- | --- | --- | --- | --- | --- | --- | --- | --- |
|  |  | AAK1 | BIKE | GAK | STK16 |  |  |  |  |
| (5Z)-7-Oxozeaenol | Investigational | 13%<sup>***</sup> | 3.8%<sup>***</sup> | 10%<sup>***</sup> | 100%<sup>***</sup> | NA | Pu et al.,2020 | ID: 20211 | TAK1 (8 nM), VEGFR2 (52 nM), ERK2 (80 nM) |
| Sunitinib | Approved (cancer) | 11 nM<sup>*</sup> | 5.5 nM<sup>*</sup> | 20 nM<sup>*</sup> | 250 nM<sup>*</sup> | NA | Davis et al., 2011 | Davis et al., 2011 | VEGFR2 (80 nM), PDGFRβ (2 nM), FLT3, KIT, PDGFRα |
| Erlotinib | Approved (cancer) | 1200 nM<sup>*</sup> | 1200 nM<sup>*</sup> | 3 nM<sup>*</sup> | >9000 nM<sup>*</sup> | 910 nM | Asquith et al., 2019 | Davis et al., 2011 | ErbB1 (0.7 nM), STK10 (19 nM), YSK4 (25 nM), SLK (26 nM) |
| Gefitinib | Approved (cancer) | >10,000 nM<sup>***^</sup> | >10,000 nM<sup>***^</sup> | 13 nM<sup>*</sup> | NA | 420 nM | Asquith et al., 2019 | Davis et al., 2011 | EGFR |
| Baricitinib | Approved (RA) | 17 nM<sup>*</sup> | 40 nM<sup>*</sup> | 134 nM<sup>*</sup> | 69 nM<sup>*</sup> | NA | Stebbing et al., 2021 | Klaeger et al., 2017 | JAK1 (5.9 nM), JAK2 (5.7 nM), ROCK1/2, TYK2 (53 nM), CAMK2A, MAP3K2, PRPF4B |
| RMC-76 | Experimental | 4 nM<sup>**</sup> | 3.8 nM<sup>**^</sup> | 280 nM<sup>**^</sup> | NA | NA | Figure S1 | NA | CDC42BPB, PRKD3, CLK1, CLK4, ULK3, MINK1 |
| RMC-242 | Experimental | 4100 nM<sup>**^</sup> | >15000 nM<sup>**^</sup> | 12 nM<sup>**</sup> | NA | 78 nM<sup>^</sup> | Martinez-Gualda et al.,2021 | NA | NA |
| SGC-GAK-1 | Experimental | NA<br>86%<sup>***</sup> | NA<br>92%<sup>***</sup> | 2 nM<sup>*</sup><br>9.7%<sup>***</sup> | NA<br>84%<sup>***</sup> | 48 nM | Asquith et al., 2019 | Asquith et al., 2019 | RIPK2 (110 nM), ADCK3 (190 nM), NLK (520 nM) |
| STK-16-1N-1 | Experimental | NA<br>13%<sup>***</sup> | NA<br>9.5%<sup>***</sup> | NA<br>46%<sup>***</sup> | 295 nM<sup>**</sup><br>0.65%<sup>***</sup> | NA | Liu et al., 2016 | Liu et al., 2016 | PI3Kδ (856 nM), PI3Kγ (867 nM) |

^ Data obtained in this study

(5Z)-7-oxozeaenol, an investigational anticancer natural product, is an ATP-competitive inhibitor (Ninomiya-Tsuji et al., 2003; Wu et al., 2013) that potently binds BIKE, GAK and AAK1 (3.8%, 10% and 13% binding of control, respectively, at 10 μM) (**Table 1**), previously demonstrating a broad-spectrum antiviral activity against DENV, Venezuelan equine encephalitis virus (VEEV/TC-83), and EBOV (Pu et al., 2020). We measured a dose-dependent inhibition of SARS-CoV-2 infection in Vero E6 and Calu-3 cells upon drug treatment with (5Z)-7-oxozeaenol with EC_50_ values of 2.3 μM, and 2.9 μM, respectively (**Figure 2B and C**). While mild toxicity was observed in Vero E6 cells with a CC_50_ of 18 μM, no apparent toxicity was measured at the tested concentrations in Calu-3 cells (**Figure 2B and C**).

Sunitinib, an approved multi-kinase inhibitor with potent binding to AAK1 (dissociation constant (K_d_=11 nM)), BIKE (K_d_=5.5 nM), and GAK (K_d_=20 nM) (**Table 1**) (Davis et al., 2011) demonstrated a moderate anti-SARS-CoV-2 effect in Vero E6 (EC_50_=2.4 μM) and Calu-3 (EC_50_=2.9 μM) cells (**Figure 2B and C**). Like (5Z)-7-oxozeaenol, sunitinib reduced cellular viability in Vero E6 cells (CC_50_=12.2 μM), yet had no apparent toxicity in Calu-3 cells (CC_50_>20 μM).

Erlotinib, an approved anticancer EGFR inhibitor (K_d_=0.7 nM), showed no anti-SARS-CoV-2 activity and no cellular toxicity in both cell lines (EC_50_>20 μM, CC_50_>20 μM) (**Figure 2B and C**). Notably, while erlotinib potently binds GAK *in vitro* (K_d_=3 nM) (Davis et al., 2011), its intracellular anti-GAK activity is low with an IC_50_ of 910 nM, as measured by a nanoluciferase-based bioluminescence resonance energy transfer (NanoBRET) assay (**Table 1**) (Asquith et al., 2019; Asquith et al., 2020). Gefitinib, another approved drug with potent GAK binding activity (K_d_=13 nM) (Davis et al., 2011) and moderate anti-GAK activity via the NanoBRET assay (IC_50_=420 nM) (**Table 1**), moderately suppressed SARS-CoV-2 infection in both cell lines (EC_50_=3.9 - 5 μM), with some toxicity in Vero E6 (CC_50_=13 μM) but not in Calu-3 cells (CC_50_>20 μM) (**Figure 2B and C**).

Baricitinib, a janus kinase (JAK) inhibitor approved for inflammatory diseases and used for COVID-19 treatment for its predicted potential effect on AAK1-mediated SARS-CoV-2 entry, beyond JAK-mediated inflammation (Richardson et al., 2020; Stebbing et al., 2020), showed weak activity against SARS-CoV-2 in Vero E6 (EC_50_=13.6 μM) and largely no antiviral activity in Calu-3 cells (EC_50_>20 μM) (**Figure 2B and C**), in line with its weaker anti-NAK activity (AAK1, K_d_=17 nM; BIKE, K_d_=40 nM; and GAK, K_d_=134 nM) (**Table 1**).

### 3.3. Sunitinib and erlotinib have a synergistic anti-SARS-CoV-2 effect

To determine whether a synergistic effect could be achieved with a combination treatment of compounds with anti-AAK1/BIKE and GAK activity, we measured the anti-SARS-CoV-2 activity of combination treatment with sunitinib and erlotinib. Calu-3 cells were pretreated for 1 hour with the combination treatment followed by infection with rSARS-CoV-2/Nluc (MOI=0.05). The effects of treatment on viral replication and cellular viability were measured at 24 hpi via Nluc and alamarBlue assays, respectively. Treatment with combinations of sunitinib and erlotinib exhibited synergistic inhibition of SARS-CoV-2 infection with a synergy volume of 38 μM^2^% at the 95% CI and calculated antagonism of −3.41 μM^2^%(**Figure 3A**) with no synergistic toxicity (**Figure 3B**). Based on Prichard et al. (Prichard and Shipman, 1996), this level of synergy is considered minor but significant. These results are in agreement with our reported *in vitro* and *in vivo* HCV, DENV and EBOV data (Neveu et al., 2015; Bekerman et al., 2017), and point to combinations of anti-NAK inhibitors as a potential anti-coronaviral strategy.

**Figure 3.**
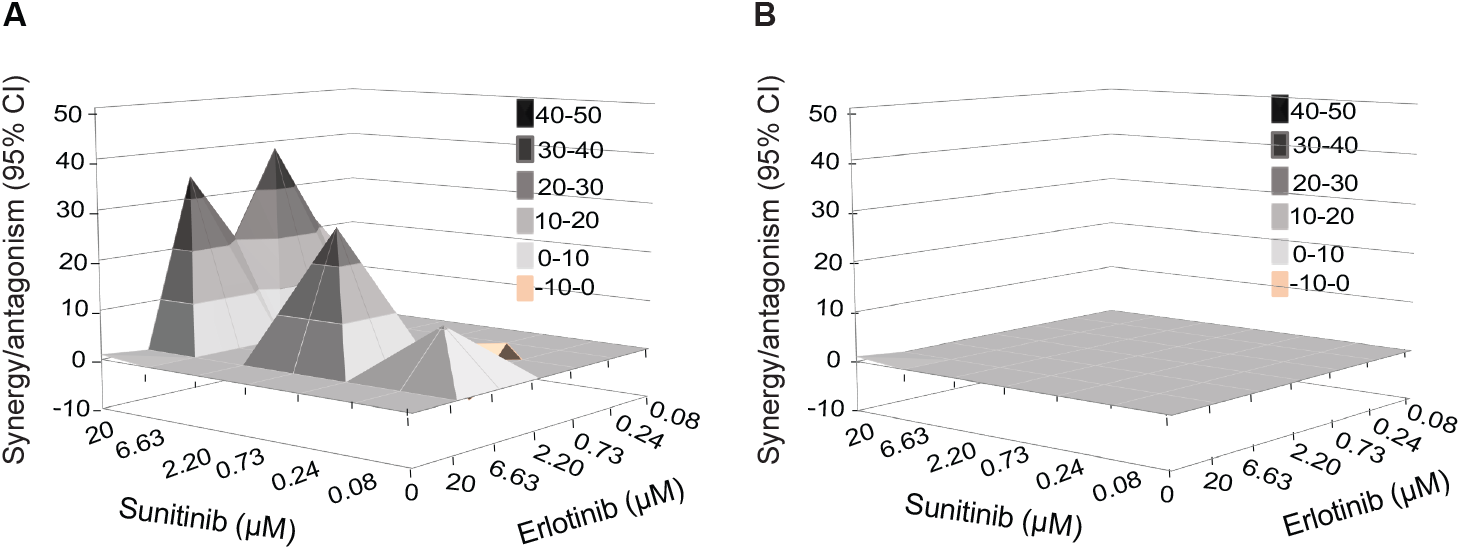
Combination treatment with sunitinib/erlotinib have synergistic effect against SARS-CoV-2. (**A** and **B**) Synergy/antagonism of sunitinib/erlotinib combination treatment on antiviral effect measured in Calu-3 cells infected with rSARS-CoV-2/Nluc (USA-WA1/2020 strain; MOI=0.05) at 24 hpi via Nluc assays (**A**) and on cellular viability measured at 24 hpi in the same samples via alamarBlue assays (**B**). Data represents the differential surface analysis at the 95% confidence interval (CI), analyzed via the MacSynergy II program. Synergy and antagonism are indicated by the peaks above and below the theoretical additive plane, respectively. The level of synergy or antagonism is depicted by the color code on the figure. Data are representative of 2 independent experiments with 3 replicates each.

### 3.4. Novel, chemically distinct NAK inhibitors potently inhibit SARS-CoV-2 infection *in vitro*

Next, we determined whether chemically distinct, novel small molecules with anti-NAK activity (**Table 1 and Figure 4A**) can suppress SARS-COV-2 infection. RMC-76 is a pyrrolo [2,3-*b*]pyridine (compound 21b in (Verdonck et al., 2019)) with potent anti-AAK1 (IC_50_=4.0 nM) and anti-BIKE (K_d_=3.8 nM) (**Table 1**) activity and reported activity against DENV and EBOV. RMC-76 effectively suppressed SARS-CoV-2 infection in both Vero E6 (EC_50_=0.3 μM) and Calu-3 (EC_50_=1 μM) cells (**Figure 4B and C**), with no apparent toxicity in both cell lines (CC50>20 μM). We further evaluated the selectivity of RMC-76 via kinome profiling in cell lysates using multiplexed inhibitor beads kinome profiling coupled with mass spectrometry (MIB/MS) (Duncan et al., 2012; Asquith et al., 2018; Asquith et al., 2019). RMC-76 showed highest selectivity towards AAK1 and BIKE, with lower affinity for the off-target kinases CDC42BPB, PRKD3, CLK1, CLK4, ULK3, and MINK1 (**Table 1 and Supplementary Figure S1**).

**Figure 4.**
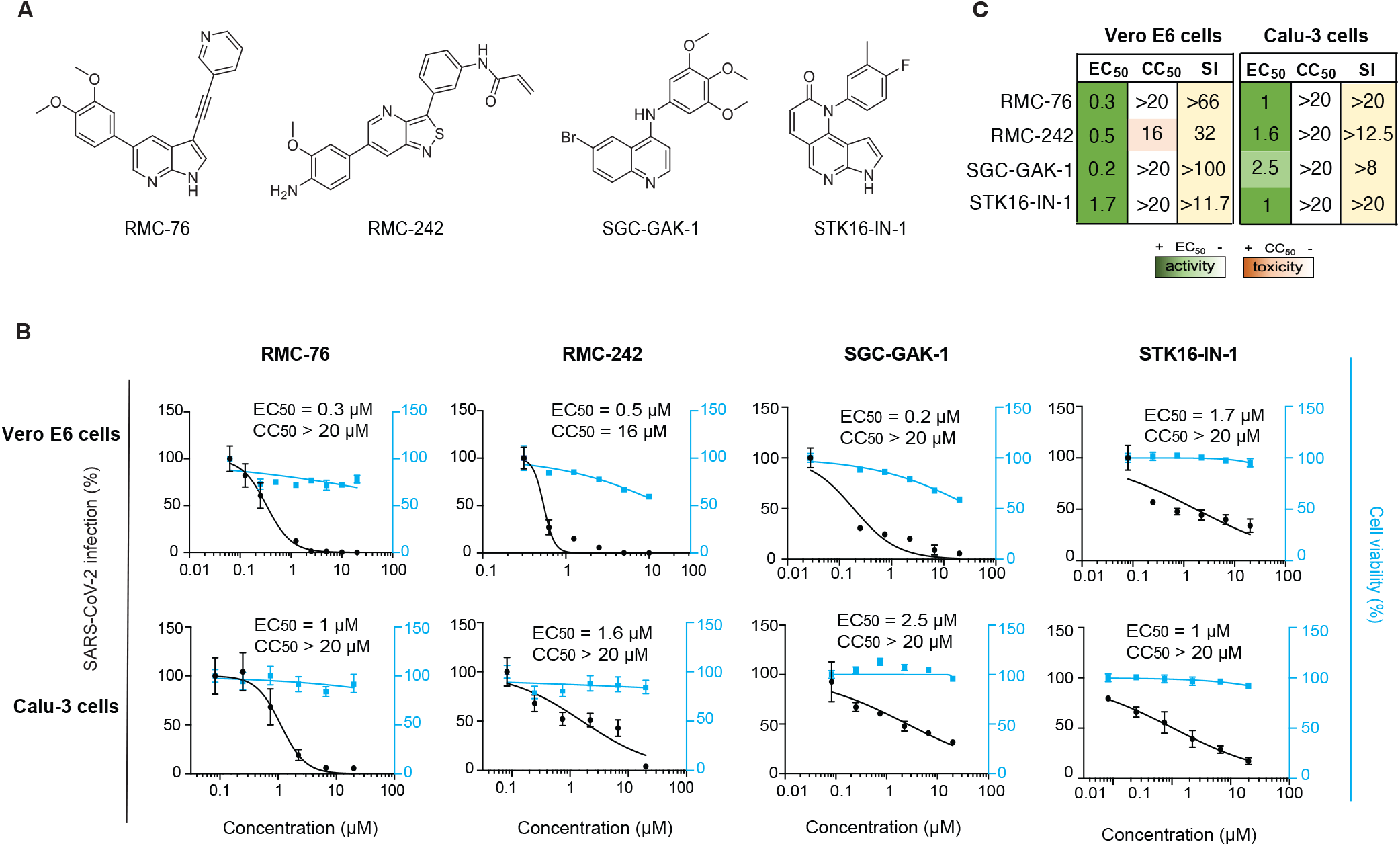
Novel compounds with anti-NAK activity suppress SARS-CoV-2 infection. (**A**) Chemical structures of RMC-76, RMC-242 and SCG-GAK-1, and STK16-IN-1. (**B**) Dose response of WT SARS-CoV-2 infection to the indicated compounds in Vero E6 cells infected with rSARS-CoV-2/Nluc and in Calu-3 cells infected with USA-WA1/2020 strain (MOIs=0.05) (black) measured via plaque and Nluc assays at 24 hpi, respectively. Dose response of cellular viability to the indicated compounds measured via alamarBlue assays are shown in blue. (**C**) Heat maps of the EC_50_ and CC_50_ values of the inhibitors color-coded based on the antiviral activity and cell viability measured in the indicated cell lines. Selectivity indices (SI, CC_50_ to EC_50_ ratios) greater than 5 are depicted in yellow. Data in all panels are representative of 2 or more independent experiments. Individual experiments had 3 or greater biological replicates. Shown are means ± SD.

RMC-242, an *N*-(3-(6-(4-amino-3-methoxyphenyl) isothiazolo [4,3-*b*]pyridin-3-yl)phenyl) acrylamide (compound 7b in (Martinez-Gualda et al., 2021)) with anti-GAK (IC_50_=12 nM) activity (**Table 1**) has previously shown anti-DENV activity. RMC-242 dose-dependently suppressed SARS-CoV-2 infection in Vero E6 (EC_50_=0.5 μM) and Calu-3 (EC_50_=1.6 μM) cells. While RMC-242 exhibited mild toxicity in Vero cells (CC_50_=16 μM), no apparent toxicity was measured in Calu-3 cells (CC_50_>20 μM) (**Figure 4B and C**). SGC-GAK-1, a 4-anilinoquinoline,(compound 11 in (Asquith et al., 2019)) with potent anti-GAK activity (K_d_=2 nM), has previously shown potent anti-DENV activity (Saul et al., 2020). SGC-GAK-1 demonstrated high or moderate anti-SARS-CoV-2 effect in Vero E6 (EC_50_=0.2 μM) and Calu-3 (EC_50_=2.5 μM) cells, respectively, with no apparent cellular toxicity (CC_50_>20 μM) (**Figure 4B and C**).

Lastly, STK16-IN-1, a pyrrolonaphthyridinone compound with high selectivity against STK16 (IC_50_=295 nM) (Liu et al., 2016) effectively suppressed SARS-CoV-2 in Vero E6 (EC_50_=1.7 μM) and Calu-3 (EC_50_=1 μM) cells with no apparent toxicity in both cell lines (CC_50_>20 μM) (**Figure 4B and C**).

These results validate NAKs as druggable antiviral targets and point to their pharmacological inhibition as a potential anti-SARS-CoV-2 strategy.

### 3.5. RMC-76 suppresses early and late stages of the SARS-CoV-2 life cycle

To pinpoint the stage of the viral life cycle that is inhibited by compounds with anti-NAK activity, we next conducted time-of-addition experiments in Calu-3 cells infected with rSARS-CoV-2/WT (MOI=1). RMC-76 (with potent anti-AAK1/BIKE activity) was added at 2, 5 or 8 hpi (**Figure 5A**). Cell culture supernatants were harvested at 10 hpi, which represents a single cycle of SARS-CoV-2 replication, and infectious viral titers were measured by plaque assay. Treatment with RMC-76 initiated upon infection onset and maintained throughout the 10-hour experiment (0 to 10) suppressed viral infection by 93%±4.1% relative to the DMSO control. RMC-76 treatment during the first 2 hours of infection only (0 to 2) suppressed viral infection by +89%±5.9% relative to DMSO, indicating that RMC-76 suppresses viral entry (**Figure 5B**). Nevertheless, addition of RMC-76 at 2 hpi (2 to 10) reduced viral infection by 77%±9.4% revealing a potential effect in a post-entry stage (**Figure 5B**). Whereas treatment with RMC-76 initiated 5 hpi (5 to 10) had no effect on viral infection, treatment initiated at 8 hpi (8 to 10) reduced SARS-CoV-2 infection by 33%±10.6%(**Figure 5B**). These results provide evidence that compounds with anti-AAK1/BIKE activity suppress viral entry and a later stage of the SARS-CoV-2 life cycle (assembly and/or egress), but likely not viral RNA replication.

**Figure 5.**
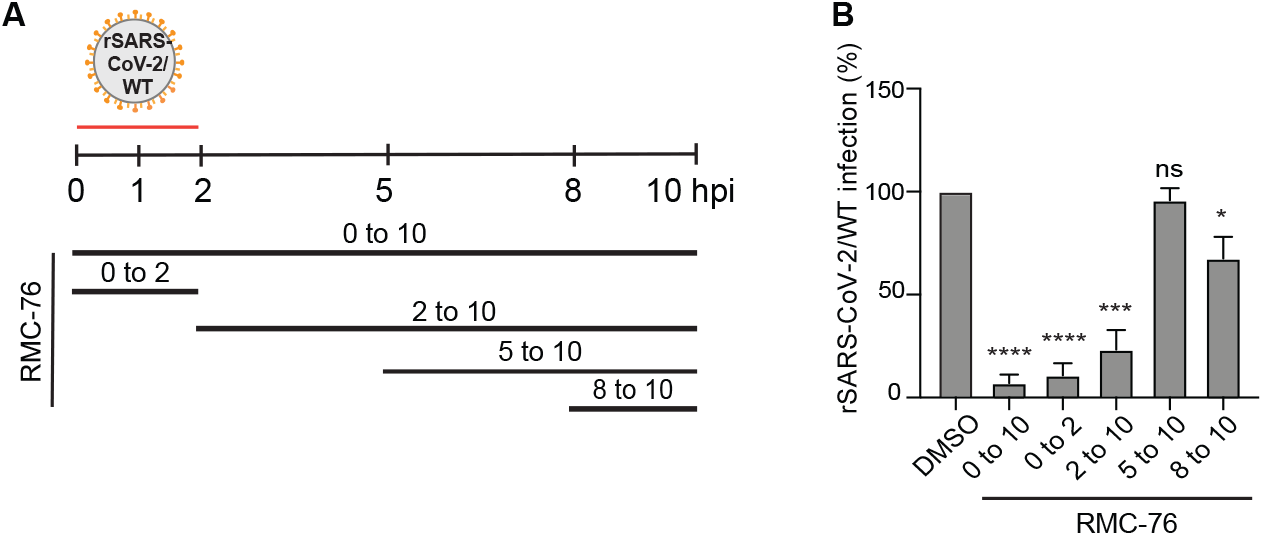
RMC-76 inhibits temporally distinct stages of SARS-CoV-2 life cycle. (**A**) Schematic of the time-of-addition experiments shown in panel **B. (B**) Calu-3 cells were infected with rSARS-CoV-2/WT at an MOI of 1. At the indicated time points, 5 μM RMC-76 or DMSO were added to the infected cells. Supernatants were collected 10 hpi and infectious virus titers were measured by plaque assays. Data are relative to DMSO and are representative of 2 independent experiments, each with 2 replicates, means ± SD are shown. *P < 0.05; ***P < 0.001; ****P < 0.0001 by 1-way ANOVA followed by Dunnett’s multiple comparisons test. Ns, non-significant.

### 3.6. NAKs are required for SARS-CoV-2 pseudovirus entry

Since RMC-76 demonstrated an effect on viral entry and SARS-CoV-2 is thought to enter target cells in part via CME (Bayati et al., 2021; Koch et al., 2021), we tested the hypothesis that NAKs are involved in mediating SARS-CoV-2 entry. To this end, we first used vesicular stomatitis virus encapsidated RNA pseudotyped with the SARS-CoV-2 spike glycoprotein expressing a luciferase reporter gene (rVSV-SARS-CoV-2-S) (Khanna et al., 2020; Saul et al., 2021). Calu-3 cells depleted of the individual NAKs (**Figure 1B**) were infected with rVSV-SARS-CoV-2-S followed by luciferase assays at 24 hpi (**Figure 6A**). Silencing of NAKs expression suppressed rVSV-SARS-CoV-2-S infection by 50-70% relative to siNT (**Figure 6B**).

**Figure 6.**
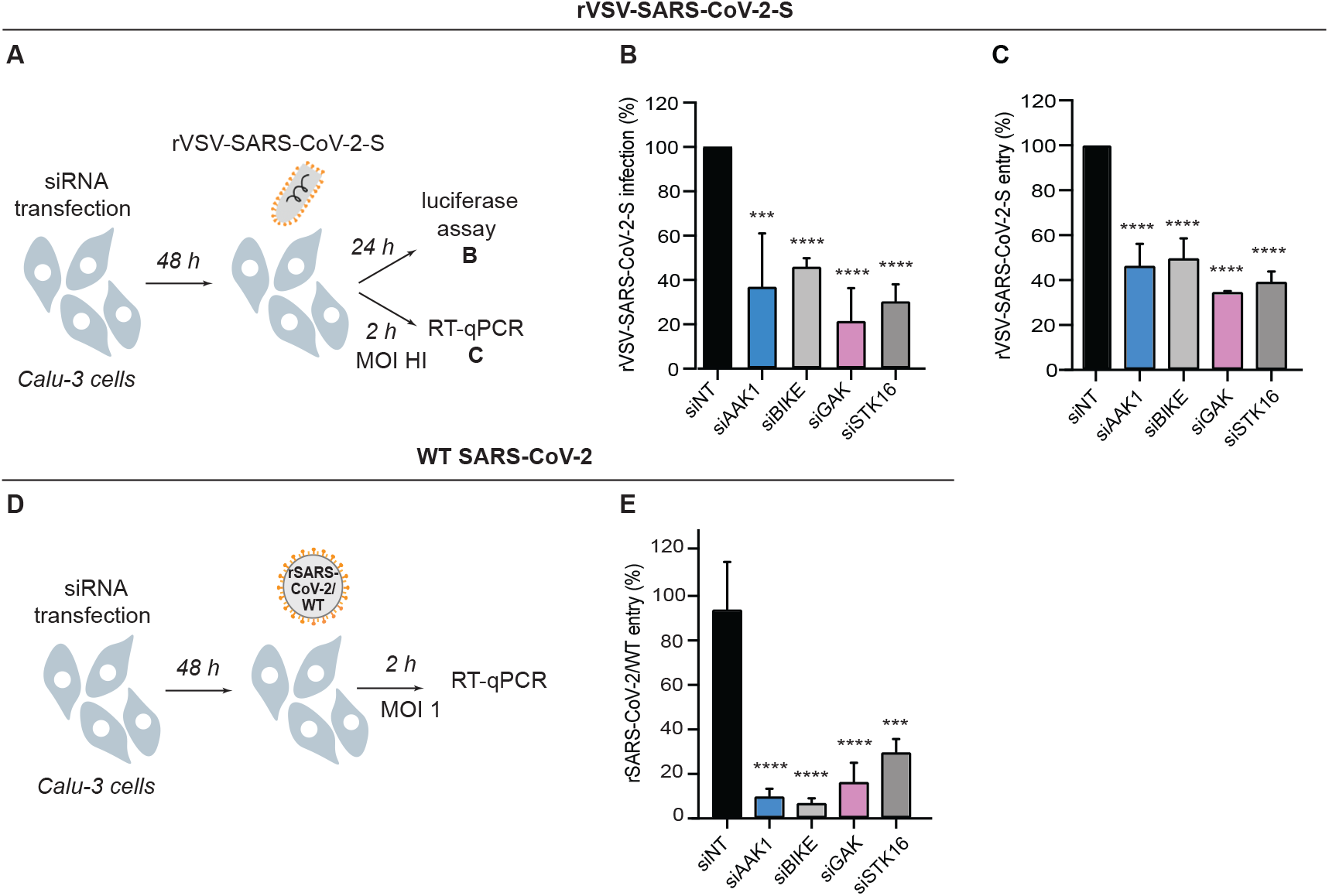
NAKs are required for the entry of pseudovirus and WT SARS-CoV-2. (**A**) Schematic of the experiments shown in panels **B** and **C**. (**B**) rVSV-SARS-CoV-2-S infection at 24 hpi of Calu-3 cells depleted of the indicated NAKs-by siRNAs (**Figure 1B**) measured via luciferase assays. (**C**) rVSV-SARS-CoV-2-S entry at 2 hpi of Calu-3 cells (high MOI) depleted of the indicated NAKs by siRNAs measured via RT-qPCR. (**D**) Schematic of the experiment shown in panel **E**. (**E**) rSARS-CoV-2/WT entry at 2 hpi of Calu-3 cells (MOI=1) depleted of the indicated NAKs by siRNAs measured via RT-qPCR. Data in all panels are representative of 2 or more independent experiments. Individual experiments had 3 biological replicates, means ± SD are shown. Data are relative to siNT (**B, C, E**). ***P < 0.001; ****P < 0.0001 by 1-way ANOVA followed by Dunnett’s multiple comparisons test.

To confirm that the observed defect was indeed in viral entry, we next quantified the level of intracellular viral RNA at 2 hpi of Calu-3 cells depleted of the individual NAKs (**Figure 1B and 6A**) with a high-inoculum of rVSV-SARS-CoV-2-S. NAKs depletion reduced the intracellular viral copy number by 50-65% relative to siNT as measured via RT-qPCR, supporting a role for NAKs in SARS-CoV-2 entry (**Figure 6C**).

### 3.7. NAKs are required for WT SARS-CoV-2 entry

To define the role of NAKs in entry of WT SARS-CoV-2, we conducted entry experiments in Calu-3 cells infected with rSARS-CoV-2/WT virus (MOI=1) (**Figure 6D**). Suppression of NAKs expression (**Figure 1B**) reduced the level of intracellular viral RNA measured via RT-qPCR at 2 hpi by 90%±3.3% (siAAK1), 93%±2.0% (siBIKE), 83%±8.5% (siGAK), and 70%±5.8% (siSTK16) relative to siNT, respectively (**Figure 6E**).

Together, these findings provide genetic evidence that the four NAKs are involved in mediating SARS-CoV-2 infection and entry.

## 4. Discussion

We have previously demonstrated that AAK1 and GAK regulate intracellular trafficking of multiple RNA viruses via phosphorylation of the AP1 and AP2 complexes and represent targets for broad-spectrum antivirals (Neveu et al., 2015; Bekerman et al., 2017; Pu et al., 2018; Xiao et al., 2018; Verdonck et al., 2019; Pu et al., 2020). We have also reported the requirement for BIKE in DENV and unrelated RNA viral infections in part via AP2M1 phosphorylation (Pu et al., 2020). Nevertheless, the roles of NAKs in coronaviral infection remained largely unknown. Moreover, to our knowledge, the role of STK16 in viral infection has not been reported to date. Here, by integrating virology, genetic and pharmacological approaches, we demonstrate a requirement for all four NAKs in SARS-CoV-2 infection. Moreover, we provide a proof of concept for the potential utility of NAK inhibition as a strategy for treating SARS-CoV-2 infection.

AAK1 was previously identified as a candidate target for SARS-CoV-2 treatment via an artificial intelligence screen (Richardson et al., 2020; Stebbing et al., 2020). Using an RNAi approach, we provide evidence that AAK1 is required for SARS-CoV-2 infection in a biologically relevant cell culture model, in agreement with a recent study demonstrating a requirement for AAK1 in other human lung cells (Puray-Chavez et al., 2021). Beyond AAK1, we provide genetic evidence that BIKE, GAK and STK16, NAKs whose functional relevance in SARS-CoV-2 has remained unknown, are also proviral factors. To our knowledge, this is the first evidence that STK16 is required for any viral infection.

We provide evidence that NAKs are involved in regulating SARS-CoV-2 entry. Depletion of NAKs reduced the entry of pseudo- and WT SARS-CoV-2 via both Nluc assays and entry-specific assays measuring viral RNA at 2 hpi. Moreover, time-of-addition experiments revealed that pharmacological inhibition of AAK1/BIKE potently inhibited viral infection when added in the first two hpi. While the precise roles of NAKs in viral entry remains to be elucidated, phosphorylation of AP2M1, NAK substrate involved in CME, was reported to be induced upon SARS-CoV-2 infection in lung cells (Puray-Chavez et al., 2021) and to be required for SARS-CoV-2 pseudovirus entry (Wang et al., 2020). Notably, a conserved tyrosine motif in the cytoplasmic tail of ACE2 was shown to mediate its interaction with AP2M1 and to be essential for SARS-CoV-2 entry into cells with low TMPRSS2 expression (Wang et al., 2020). Sorting into late endosomes is the main penetration route of SARS-CoV-2 into cells with low or no TMPRSS2 expression. In TMPRSS2 expressing cells, beyond fusion-mediated penetration at the plasma membrane, SARS CoV 2 sorting into the endocytic pathway also appears to play a role (Bayati et al., 2021; Koch et al., 2021). Our findings that NAKs, regulators in the endocytic pathway, are required for effective entry of WT SARS-CoV-2 into Calu-3 (high TMPRSS2 expressing) cells support the latter.

Beyond viral entry, the time-of-addition experiments provide evidence that NAKs are also involved in regulating later stage(s) of the viral life cycle, likely assembly and/or egress, but not viral RNA replication. This is in agreement with our prior findings involving their roles in unrelated RNA viral infections such as DENV, EBOV, and VEEV (Neveu et al., 2015; Bekerman et al., 2017; Pu et al., 2020; Saul et al., 2020; Huang et al., 2021; Saul et al., 2021; Saul et al., 2021). Together, these findings provide evidence that NAKs regulate temporally distinct stages of the SARS-CoV-2 life cycle and represent candidate targets for anti-SARS-CoV-2 treatment.

Using compounds with potent anti-NAK activity as pharmacological tools we provide further support for NAKs requirement in SARS-CoV-2 infection and validate them as antiviral targets in lung epithelial cells. RMC-76 and sunitinib, novel and approved compounds with potent anti-AAK1/BIKE activity demonstrated antiviral effect against SARS-CoV-2. This is in line with a previous study reporting suppression of SARS-CoV-2 infection with a selective AAK1 inhibitor (SGC-AAK1-1) (Puray-Chavez et al., 2021). The role of AAK1 and BIKE in the entry of SARS-CoV-2, a stage of the viral life cycle that is inhibited by RMC-76, supports a hypothesis that inhibition of NAKs, at least in part, mediates the antiviral effect of this compound.

Compounds with anti-GAK activity have demonstrated variable antiviral effect. Erlotinib demonstrated no antiviral effect when used individually, gefitinib showed moderate antiviral activity, RMC-242 and SGC-GAK-1 showed potent or moderate antiviral activity against SARS-CoV-2, respectively. While all four compounds potently bind GAK in an *in vitro* binding assay (K_d_ values of: 3 nM (erlotinib), 13 nM (gefitinib), 12 nM (RMC-242), and 2 nM (SGC-GAK-1)), their anti-GAK activity in the cell-based target engagement nanoBRET assay (the most relevant biochemical measurement for predicting a biological effect in cells) is variable (IC_50_ values of: 910 nM (erlotinib), 420 nM (gefitinib), 78 nM (RMC-242), 48 nM (SGC-GAK-1)), yet it correlates with their anti-SARS-CoV-2 effects. The STK16 inhibitor, STK16-IN-1 demonstrated anti-SARS-CoV-2 effect. Combined with the siRNA data, this finding validates the requirement for STK16 in SARS-CoV-2 infection.

Baricitinib, an approved anti-inflammatory drug, only weakly suppressed SARS-CoV-2 infection in both Vero E6 and Calu-3 cells, in agreement with previous reports (Wang et al., 2020; Stebbing et al., 2021). Baricitinib shows moderate binding to AAK1 (K_d_=17 nM) and BIKE (K_d_=40 nM) and low binding to GAK (K_d_=134 nM) and STK16 (K_d_=69 nM). Nevertheless, its anti-NAKs activity in a cell-based target engagement assay is not known, and it was shown to inhibit AP2M1 phosphorylation only at a high concentration (10 μM) (Wang et al., 2020). Additionally, the kinome of baricitinib reveals very potent binding to diverse kinases, with ROCK1/2, TYK2, CAMK2A, MAP3K2, and PRPF4B, being just several examples (Klaeger et al., 2017). As with other multi-kinase inhibitors, it is difficult to predict the antiviral effect of baricitinib, as it is driven by its net effect on both proviral and antiviral targets, as we have previously shown with erlotinib and sunitinib (Bekerman et al., 2017). Baricitinib is studied in COVID-19 patients (Kalil et al., 2021; Marconi et al., 2021) for its anti-inflammatory effect mediated by its known target, JAK, and its NAK-mediated antiviral effect predicted by an artificial intelligent screen (Richardson et al., 2020; Stebbing et al., 2020). Our findings suggest that the benefit demonstrated with baricitinib clinically likely does not result from its predicted anti-NAK effect and point out a limitation of the artificial intelligence approach for drug discovery in predicting antiviral activity.

Lastly, we show that combining compounds with anti-AAK1/BIKE and anti-GAK activities may provide a synergistic antiviral effect against SARS-CoV-2 *in vitro*. This finding is in agreement with our prior data with sunitinib/erlotinib combinations in HCV, DENV, and EBOV *in vitro* (Neveu et al., 2015; Bekerman et al., 2017) and the finding that their combination protected 85% and 50% of mice from DENV and EBOV challenges, respectively (Bekerman et al., 2017; Pu et al., 2018). AAK1/BIKE and GAK have partially overlapping functions (Zhang et al., 2005; Neveu et al., 2015; Pu et al., 2020), which may explain moderate or no antiviral effect with drug alone, yet synergistic activity upon treatment with sunitinib/erlotinib combinations. The observed synergistic effect may also result from inhibition of additional targets by these drugs.

In summary, these findings validate NAKs as candidate druggable targets for antiviral therapy against SARS-CoV-2 infection and provide a proof of concept that anti-NAKs approaches may have a utility for treating SARS-CoV-2 and possibly other coronavirus infections.

## Figure Legends

**Supplementary Figure S1. RMC-76 is highly selective and highly potent for AAK1 and BIKE**.

SUM159 cell lysates were incubated with DMSO or 1 μM RMC-76 for 30 min on ice. Kinases were then affinity purified using multiplexed inhibitor beads (MIBs) and analyzed by mass spectrometry. Kinase abundance was quantified label-free using MaxQuant software. Bars represent the ratio of label-free quantification values for the indicated kinase in lysates treated with drug over DMSO control lysates. Data means ± SD of 2 replicates.

## Acknowledgements and funding

This work was supported by awards number W81XWH-16-1-0691 and W81XWH2110456 from the Department of Defense, Congressionally Directed Medical Research Programs (CDMRP), award number GRANT12393481 from The Defense Threat Reduction Agency (DTRA) Fundamental Research to Counter Weapons of Mass Destruction, and award number RO1-AI158569–01 from the National Institute of Allergy and Infectious Diseases (NIAID) to S.E. This work was partly supported by the NIH Common Fund Illuminating the Druggable Genome (IDG) program (NIH Grant U24DK116204). S.E. is a Chan Zuckerberg Biohub investigator. M.K. was supported by a Postdoctoral Fellowship in Translational Medicine by the PhRMA Foundation. We thank the Stanford *in vitro* BSL3 service Center and its Director Dr. Jaishree Garhyan for assistance in the BSL3. We also thank the staff of Stanford Clinical Virology Laboratory for their help sequencing the SARS-CoV-2 USA-WA1/2020 isolate used in this work.

## Author Contributions

M.K., and S.E., wrote the first version of the manuscript. M.K., S.S., L.G., M.K.S., N.B., M.P.E., and G.J., designed and performed the experiments and conducted data analysis. C.Y., J.G.P., B.M.G., and J.J., provided reagents and guidance. S.E., B.A.P., L.M.S., C.R.M.A., A.N., and S.D., provided scientific oversight and guidance. S.E., provided the funding for this study.

