## Supplementary figures and images for "Numb-associated kinases are required for SARS-CoV-2 infection and are cellular targets for therapy"

### Supplementary Figure S1

#### Supplementary Figure S1

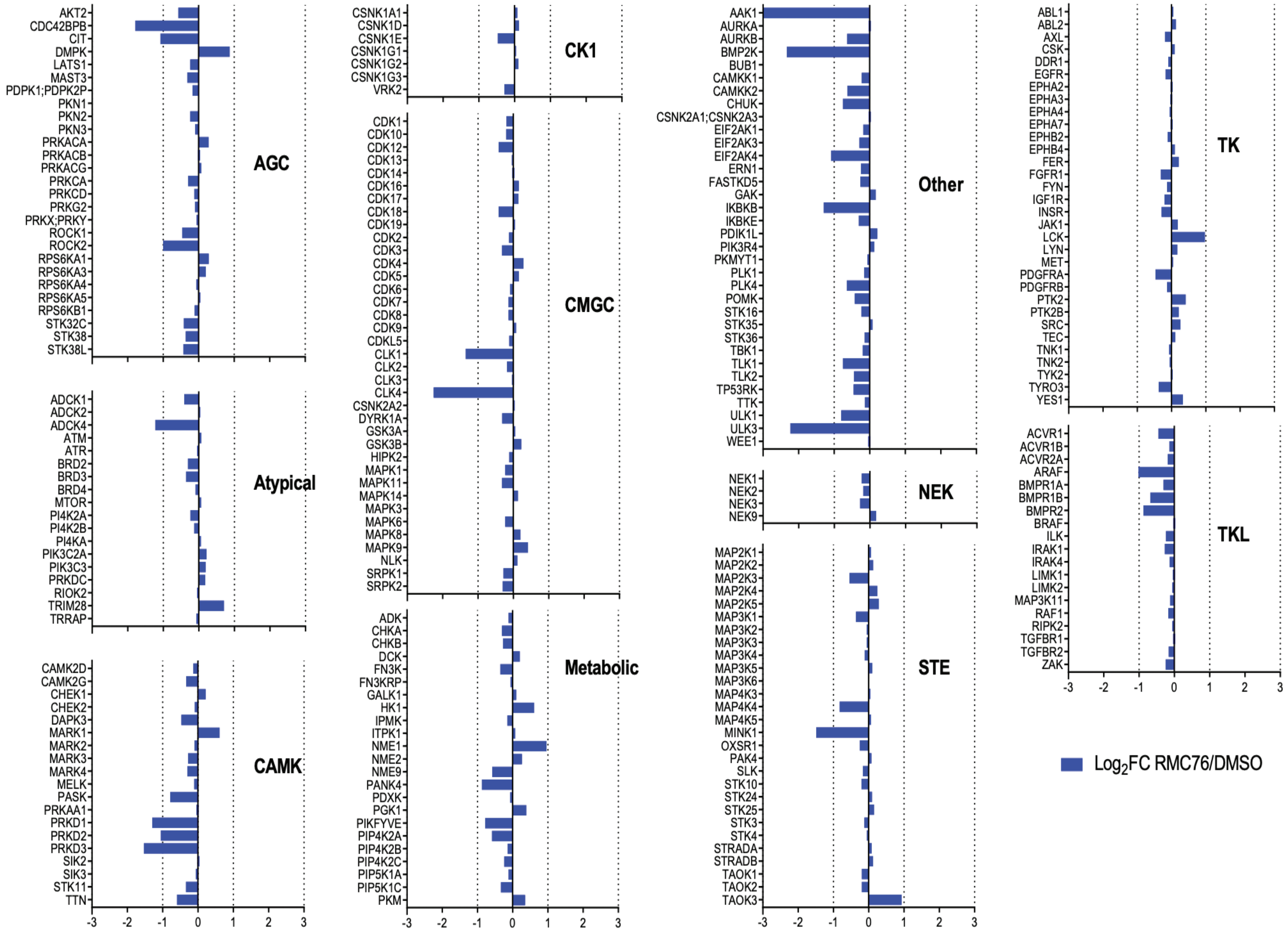
